# The equivocal mean age of parents in a cohort

**DOI:** 10.1101/371245

**Authors:** François Bienvenu

## Abstract

The mean age at childbirth is an important notion in demography, ecology and evolution, where it is used as a measure of generation time. A standard way to quantify it is to compute the mean age of the parents of all offspring produced by a cohort, and the resulting measure is thought to represent the mean age at which a typical parent produces offspring. In this note, I explain why this interpretation is problematic. I also introduce a new measure of the mean age at reproduction and show that it can be very different from the mean age of parents of offspring of a cohort. In particular, the mean age of parents of offspring of a cohort systematically overestimates the mean age at reproduction, and can even be greater than the expected lifespan of parents.

## 1 Introduction

The mean age at reproduction is a central notion in the study of the evolution of reproductive timing and of the slow-fast continuum, and it also plays an important role in demography. However, as with many descriptors of populations, it is not clear how it should be defined – let alone quantified in practice. A standard way to measure it is to use the *mean age of parents of offspring produced by a cohort*: consider all offspring produced by a cohort of newborns over its lifetime; for each of these offspring, record the age that their parents (mother, in the case of a female-based model) had when the offspring was born, and take the average of these ages.

This quantity, which is is also frequently referred to as the *cohort generation time*, is straightforward to compute from complete census data. In practice however, it is usually estimated from life-tables using the following formula:

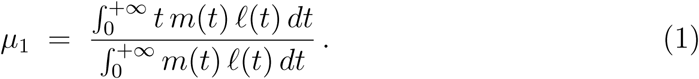

In this expression, the *survivorship function 𝓁* gives the probability that an individual of the chosen cohort reaches age *t*, and the *age-specific fertility m* represents its rate of offspring production in such a way that, assuming the individual remains alive between ages *a* and *b*, the expected number of offspring it will produce in that interval of time is 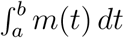. There is also a discrete-time version of formula (1):

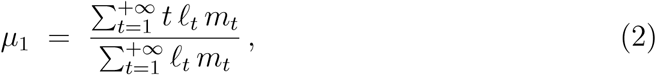

where *𝓁*_*t*_ is the probability that an individual survives to age *t* and *m*_*t*_ is the expected number of offspring produced at age *t* by individuals who reach that age.

Formulas (1) and (2) go back a long way, and are ubiquitous in the literature. They have been popularized by classic references such as Keyfitz (1968) and Coale (1972) in demography, and Charlesworth (1994) and Caswell (2001) in biology. They can also be found in more recent works of reference – including but not limited to Jørgensen and Fath (2008), Rockwood (2015) and Kliman (2016).

A consensual interpretation of *μ*_1_ is that it represents the mean age at which a typical parent produces offspring. The aim of this note is to show that this interpretation is inaccurate and can be problematic in practice. To do so, I introduce a more direct measure of the mean age at reproduction of a typical parent: consider a typical parent, and compute the average of the ages at which it gives birth to its offspring. The expected value of this average is what we term the *mean age at reproduction*. Under standard assumptions, it is given by

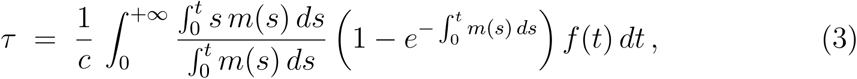

where *f* denotes the probability density function of the lifespan of an individual and the constant

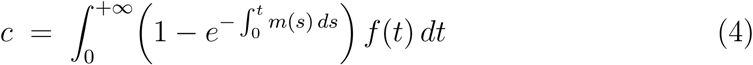

is the fraction of individuals that produce offspring during their lifetime. As with *μ*_1_, there is a discrete-time formula for *τ*:

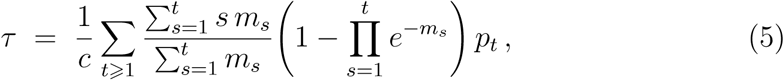

where *p*_*t*_ = ℙ(*T*_*i*_ = *t*) = *𝓁*_*t*_ − *𝓁*_*t*+1_ and

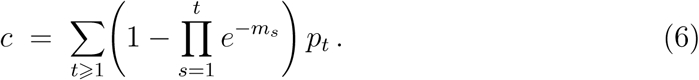

Using the expressions of *μ*_1_ and of *τ*, we show by means of concrete examples that these two quantities can differ greatly, both under simple modelling hypotheses and in real-world applications. We also investigate the relation between them mathematically. In particular, we show that *μ*_1_ is always greater than *τ*, and that the difference between the two can be arbitrarily large.

## 2 Interpretation of the expressions of *μ*_1_ and *τ*

The detailed derivations of the expressions of *μ*_1_ and *τ* are somewhat technical. They can be found in the Appendix, and a brief overview of some of the mathematical notions on which they rely is provided in the Online Supplements. Here we simply present the assumptions behind the formulas and explain what the quantities *μ*_1_ and *τ* correspond to.

Let us start with our working hypotheses. Although seldom made explicit in sources presenting *μ*_1_, the following mathematical assumptions are essential to the expressions of *μ*_1_ given in the introduction.

In the continuous-time setting, we assume (1) that the lifetimes of individuals are independent copies of a random variable *T* such that ℙ(*T* ⩾ *t*) = *𝓁*(*t*); and (2) that births are punctual random events that occur while individuals are alive (but are independent of everything else), and that there exists a function *m* such that the expected number of offspring produced by any individual alive between ages *a* and *b* is 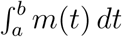 – in other words, that the birth events are the points of a point process with intensity *m*. Such models are known a Crump-Mode-Jagers processes (Crump and Mode, 1968, 1969; Jagers, 1969) – also sometimes referred to as generalized branching processes.

In the discrete-time setting, (2) is replaced by the assumption that at each age *t* = 1, 2, *…* at which it is alive, an individual produces a random number of offspring that is independent of everything else and has mean *m*_*t*_.

Under these hypotheses, it can be shown (see Section B of the Appendix) that, if we let *N* be the random variable corresponding to the number of offspring produced by a typical individual over its lifetime and *S* be the sum of the ages at which it produces them, then the quantity *μ*_1_ given in formulas (1) and (2) can rigorously be interpreted as

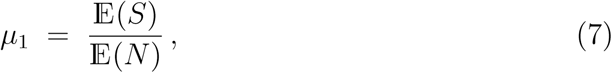

where 𝔼(*·*) denotes the expected value. This however is neither the average age at which the individual produces offspring (as often claimed) nor the mean age of the parents of the offspring produced by a cohort. Nevertheless, using the assumption that individuals are independent, by a simple application of the law of large numbers we can show (see again Section B of the appendix) that the average of the ages of the parents of the offspring produced by a cohort tends to *μ*_1_ as the size of the cohort goes to infinity.

Since the average age at which a typical individual produces offspring is the random variable *S/N*, a more natural measure for the mean age at reproduction than the ratio of expected values 𝔼(*S*)*/* 𝔼(*N*) would be the expected value of *S/N*. A minor difficulty however is that this average is well-defined only when the individual produces some offspring, i.e. when *N >* 0. Thus, we define *τ* to be the conditional expectation

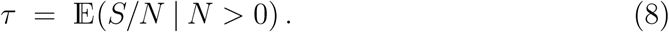

Equivalently, *τ* can be defined as follows: consider a typical parent (i.e. sample an individual uniformly at random among all individuals that produce some off-spring), and denote by *Ñ* its number of offspring and by 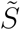 the sum of the ages at which it produces them. Then,

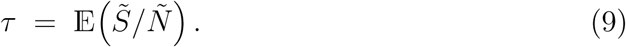

Assuming that the point processes describing birth events are Poisson point processes (or, in the discrete-time setting, that the number of offspring produced each year by each individual follows a Poisson distribution), we prove in Section C of the Appendix that *τ* is given by formula (3) (resp. (5) in discrete time). Although these formulas rely on an extra assumption compared to the expressions of *μ*_1_, observe that while the interpretation of *μ*_1_ as an average on a large cohort depends crucially on the independence of individuals, this hypothesis is not required when working with *τ*, which is truly a characteristic of individuals.

When studying evolution, it is not uncommon for one to be interested in the average of a function *z* of the ages at which a parent produces offspring rather than in the average of the ages themselves ^1^. One can then use that, letting *A* denote a uniformly chosen age at which a typical parent produces offspring, for every function *z*,

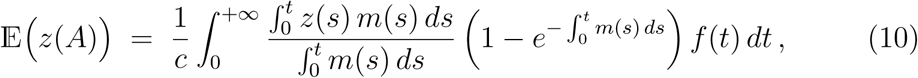

with the constant *c* given in equation (4).

Finally, let us mention that expressions of *μ*_1_ are also available for more general population structures. For instance, in matrix population models, if we let **S** be the survival matrix and **F** be the fertility matrix (i.e. if we decompose the projection matrix **A** into **A** = **S** + **F** to separate survival probabilities from fertilities) and denote by **w** the stable distribution of the population (the dominant right-eigenvector of **A**) and **e** = (1,…, 1) the row vector consisting only of ones, then we can use the following modern version of the classic formula of Cochran and Ellner (1992), which can be found in Ellner (2018):

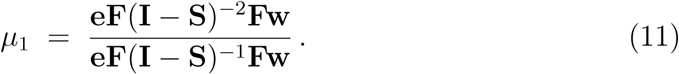

Note that (**I** - **S**)^*-*1^ = Σ_*t*⩾1_ **S**^*t-*1^ and that (**I** - **S**)^*-*2^ = Σ_*t*⩾1_ **S**^*t-*1^, so that this expression closely parallels (2). Also, the entries of **e** represent the weight given to each type of offspring when computing the average age of the parents. Taking **e** = (1,…, 1) thus corresponds the natural choice to weight all types of offspring equally. But, should we wish to give more importance to some offspring type, any vector with positive entries could be used in place of **e** – and in fact Cochran and Ellner (1992) suggest using the reproductive values as weights. See Steiner et al. (2014) and Ellner (2018) for more on this.

Unfortunately, as explained at the end of Section C of the Appendix, we have not been able to obtain an analogue of (11) for *τ*. Nevertheless, its definition as the mean age at which a typical parent produces offspring still applies in the context of matrix population models, and it is straightforward to estimate it numerically via individual-based simulations.

## 3 Examples

Let us start with a simple but fundamental example, where individuals reproduce at constant rate *m*. In that case,

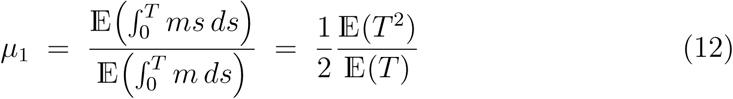

and

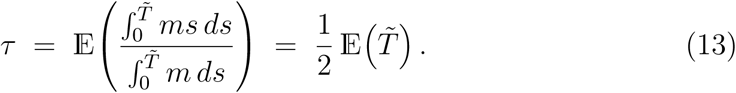

In particular, the expression of *τ* was foreseeable, because birth events are uniformly distributed on the lifetime of individuals, so on average they occur in the middle of them. Also, since

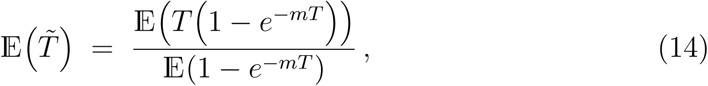

and that for all *t >* 0, 1 *-e*^*-mt*^ increases to 1 as *m* goes to infinity, it follows by monotone convergence that

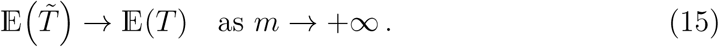

By a similar argument that can be found in Section S3 of the Online Supplements, we also have

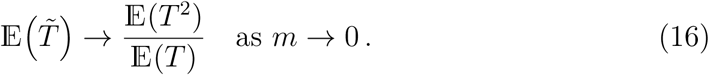

Furthermore, since 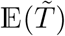 can be shown to be a decreasing function of *m* (e.g., by a standard coupling argument), we conclude that when individual reproduce at a constant rate,

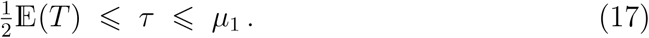

In fact, the second inequality holds for general age-specific fertility functions, as shown in Section S4 of the Online Supplements.

To make this example more concrete, let us assume that individuals die at constant rate *η*, so that *T* is an exponential variable and that *𝓁*(*t*) = *e*^*-ηt*^. In that case, after simplifications we get

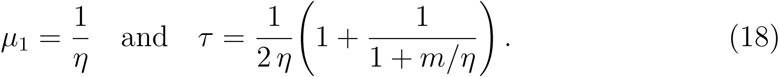

Note that here *μ*_1_ is also equal to the expected lifespan in the population. Interpreting it as the mean age at which parents reproduce would therefore lead to a contradiction, because – in the case where the fertility *m* is large enough, so that most individuals get to reproduce during their lifetime and that the lifespan of a typical parent is not very different from that of a typical individual – this would imply that, on average, the age at which an individual reproduces is the same as the age at which it dies. This is absurd, because unless individuals reproduce exactly when they die, the former has to be smaller than the latter.

We also see that for *m/η* large enough, *μ*_1_ *≈* 2*τ*. For *m* = *η*, which corresponds to the minimum ratio *m/η* to have a population that does not go extinct, the difference is already 33% of the value of *τ*.

Now consider the closely related discrete-time model where individual survive from one year to the other with probability *p* and produce Poisson(*m*) offspring at each age *t* ⩾ 1, so that

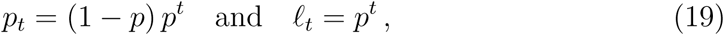

After straightforward calculations, we find that the numerator in formula (2), which corresponds to the mean sum of the ages at childbirth, is *mp/*(1 - *p*)^2^ and that the denominator is *R*_0_ = *mp/*(1 - *p*). As a result,

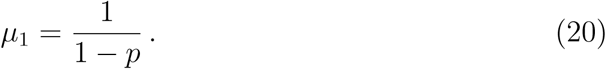

Note that this model can also be seen as a 1 × 1 matrix population model with survival matrix **S** = (*p*) and fertility matrix **F** = (*m*), so formula (11) can also be used to get this result.

Because 𝔼(*T*) = *p/*(1 − *p*), we see that

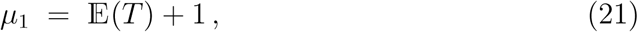

which also corresponds to the expected lifespan of individuals that reach age 1. For the same reason as before, this implies that *μ*_1_ is not credible as an estimate for the mean age at which a typical parent produces offspring.

After standard calculations that have been relegated to Section S5 of the Online Supplements, we find that

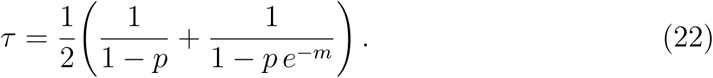

As previously, 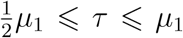, but the difference between *μ*_1_ and *τ* can be quite high, even for very reasonable values of *p* and *m*: for instance, with *p* = 0.5 and *m* = 2 the discrepancy is 30% of the value of *τ*; for *p* = 0.9 and *m* = 2, which yields a model that is similar to many real-world matrix populations models, it is already 80%.

What about real-world situations? Consider the matrix population models for the tropical palm *Astrocaryum mexicanum* (Pinero et al., 1984; Cochran and Ellner, 1992), whose projection matrices can be found in Section S6 of the Online Supplements. These models have frequently been used as examples of matrix population models, as illustrated by the fact that one of them is shipped with the ULM software for studying population dynamics (Legendre and Clobert, 1995).

Table 1 lists some relevant descriptors of these models. As already mentioned, we do not know how to compute *τ* for general matrix population models but it is straightforward to estimate it from individual-based simulations, which is what we have done here.

**Table 1:** Comparison of several measures of reproductive timing for three real-world models for the demography of the tropical palm *Astrocaryum mexicanum*, in part taken from Table 4 of Cochran and Ellner (1992): 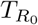 denotes the *R*_0_ generation time, which corresponds to the time it takes for the population to grow by a factor of its net reproductive rate; *Ā* is the mean age of parents of offspring in a population that has reached its stable distribution; *μ*_1_ is computed as in formula (11); 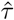 and 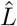 are estimates of the mean age at reproduction and of the expected lifespan conditional on producing offspring, respectively, that were obtained using individual-based simulations (see Section S6 of the Online Supplements). All values are expressed in years.

|  | High density | Medium density | Low density |
| --- | --- | --- | --- |
| $T_{R_0}$ | 197.5 | 169.2 | 153.6 |
| $\bar{A}$ | 152.5 | 122.8 | 105.1 |
| $\mu_1$ | 275.2 | 261.8 | 275.5 |
| $\hat{\tau}$ | 152.0 | 131.3 | 184.8 |
| $\hat{L}$ | 232.2 | 207.6 | 296.0 |

The difference between *μ*_1_ and *τ* is significant in all three scenarios – a factor 2 in the *Medium density* model. Moreover, *μ*_1_ is also consistently much greater than *Ā*, the mean age of the parents of the offspring produced by the population when it has reached its stable distribution, and than *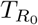*, the *R*_0_ generation time. This is intriguing, because these three quantities are all supposed to give a measure of generation time. Overall, *τ* seems to be in much better agreement with 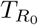 and *Ā* than *μ*_1_. Finally, and most surprisingly, we see that in the *High density* case, *μ*_1_ is greater than the expected lifespan conditional on reproduction. This counter-intuitive fact casts serious doubts on the relevance of *μ*_1_ as a measure of reproductive timing in the life-cycle.

## 4 Discussion

The expressions of *μ*_1_ and *τ* and the examples of the previous section make it clear that the mean age of the parents of the offspring produced by a cohort and the mean age at reproduction are two genuinely different notions. It is thus natural to wonder why they have not been recognized as such before. Part of the answer probably comes from the fact that precise definitions of these quantities are seldom given. For instance, in the references given above – which are or have been among the most influential in the field – *μ*_1_ is variously described as the “mean age at childbearing in the stationary population ^2^” by Keyfitz (1968); as the “mean age of childbearing in a cohort” by Coale (1972, eq. (2.10) p. 19); as the “mean age at reproduction of a cohort of females” by Charlesworth (1994, eq. (1.47a) p. 30); and as the “mean age of the mothers at the time of their daughter’s birth” by Rockwood (2015, eq. (4.12) p. 98). Yet these four definitions fail to mention explicitly the crucial fact that each offspring produced by the cohort should be considered when computing the average. In particular, they are all compatible with the idea of sampling an individual uniformly at random among the parents of a cohort and computing the mean age at which it produces offspring – and thus apply just as well to *τ*.

It is not obvious from the definitions of *μ*_1_ and *τ* how these two quantities are related – or indeed why they should differ at all. One helpful way to think about it is the following: *μ*_1_ can be seen as an *offspring-centric* measure of the mean age of parents, whereas *τ* is a *parent-centric* measure of it. Indeed, to compute *μ*_1_ we ask each newborn produced by a cohort “how old is your parent?”, while for *τ* we ask a parent “how old are you going to be when you have children?” These questions have distinct answers because they correspond to two different ways to sample a parent.

Among other things, this explains why *μ*_1_ is greater than *τ*: indeed, parents that live longer tend to have more offspring, and thus have a higher probability of being sampled via their offspring than when the sampling is done uniformly at random. As a result, they contribute more to *μ*_1_ than to *τ*. Since these parents with longer lifespans are also those that tend to have a higher mean age at reproduction, this biases *μ*_1_ upward compared to *τ*.

This also explains why the difference *μ*_1_ - *τ* goes to zero as the fertility becomes vanishingly small: in that case, the proportion of parents that give birth to more than one offspring during their lifetime goes to zero, and as the result the two parent-sampling schemes become equivalent.

Now that we better understand the link between *μ*_1_ and *τ*, two important questions arise: (1) does the difference matter in practice? (2) which descriptor should be used in which context?

There are strong arguments in favor of a positive answer to the first question. First, observe that, from a purely mathematical point of view, the difference between *μ*_1_ and *τ* can be made arbitrarily large. Indeed, recall that, when individuals reproduce at a constant rate *m, μ*_1_ = 𝔼(*T*^2^)*/* 𝔼(*T*) and 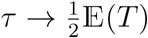 as *m* → + ∞. Thus, by choosing an appropriate distribution for the lifespan *T* and taking *m* large enough, we can make *μ*_1_ arbitrarily large and *τ* arbitrarily small. Although this requires using biologically unrealistic lifespans with large variances, this is informative because it shows conclusively that *μ*_1_ and *τ* are not just two slightly different formalizations of the same concept. Second, the examples of Section 3 show that, both under very standard modelling hypotheses and in real-world applications, *μ*_1_ can easily be roughly twice as large as *τ*. Of course, this is does not mean that the difference is going to be significant in typical real-world situations – and, in fact, since *μ*_1_ = *τ* when the lifespan of individuals is deterministic, it seems likely that *μ*_1_ and *τ* will be similar in situations where there is little variation in the lifespan of individuals – but this at least indicates that special caution should be taken when interpreting *μ*_1_.

Regarding the second question, is still unclear which of *μ*_1_ or *τ* should be favored in which context. From a practical point of view, the expressions of *τ* are, admittedly, more complex than those of *μ*_1_. This of course is not a problem for real-world applications, where they are going to be evaluated numerically; but for theoretical applications this does make exact calculations harder, if possible at all.

Another thing to take into account is the slightly different domain of validity of both measures. While the interpretation of *μ*_1_ hinges on the assumption that there are no interactions between individuals, the expression of *τ* relies on that of Poissonian births. One might cynically argue that this is hardly a problem, because both hypotheses are often used jointly in theoretical models – and never met in real-world applications. Nevertheless, there is a real difference here that should be taken into account when deciding which measure to choose.

Lastly, *τ* has the advantage of having a more direct interpretation than *μ*_1_ – and judging from the phrasing used by several authors, it seems that it is sometimes *τ* they have in mind, even when working with *μ*_1_. Moreover, the interpretation of *μ*_1_ might not be as intuitive as we usually assume; notably, the fact that it can be greater not only than the expected lifespan but also than the expected lifespan conditional on reproduction (as illustrated by the *Medium density* scenario for *Astrocaryum mexicanum* in Table 1) is likely to come as a surprise to many researchers.

## Acknowledgments

Mauricio González-Forero and Stéphane Legendre encouraged me to write this note and provided helpful comments on a first version of it. Jean-Jil Duchamps did a careful cross-checking of the math and helped me clarify a few points. Florence Débarre helped me increase the accessibility of the manuscript. Finally, my deepest thanks go to Amaury Lambert and Stephen P. Ellner who, in addition to being attentive to details, pointed me to the correct interpretation of the classic formulas for the mean age at childbirth in a cohort. Stephen Ellner should also be credited for the simple proof of Proposition 1.

## A An explicit model for the population

Here we recall and further detail the assumptions on which the expressions of *μ*_1_ and *τ* and their interpretations rely, and introduce some notation.

The setting that we use is that of a Crump-Mode-Jagers process (Crump and Mode, 1968, 1969; Jagers, 1969), where the population consists of a discrete set of individuals such that:

i. Each individual *i* has a random lifespan *T*_*i*_ with distribution *ν* and which is independent of everything else.
ii. Individual *i* produces a new offspring at age *t* for every point of *P*_*i*_ at *t* such that *t* ⩽ *T*_*i*_, where *P*_*i*_ is a point process with intensity *m* on [0, + ∞ [that is independent of everything else.

Note that the point processes *P*_*i*_ are not homogeneous (*m* is a function of the age of individuals) and that they do not have to be simple (an individual can give birth to several offspring simultaneously). For mathematical tractability however, it is often convenient to work with *Poisson point processes*. As explained in Section S1 of the Online Supplements, where a few useful results about Poisson point processes can also be found, these allow to formalize the familiar idea that events “occur at rate *m*”. While the assumption that *P*_*i*_ are Poisson point processes is not needed in the study of *μ*_1_, it will be required to derive explicit formulas for *τ*.

In this setting, the definition and interpretation of the survivorship function and of the age-specific fertility are straightforward. The survivorship is defined by ^3^

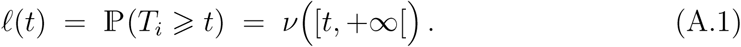

Working with the measure *ν* is convenient because it makes it possible to treat the case where *T*_*i*_ is a continuous random variable and the case where it is a discrete random variable simultaneously. However, in many applications *T*_*i*_ will have a density *f*. Thus, we will do most of our calculations with *ν* but express our final results in terms of *f* or *𝓁*, as in formulas (1) and (3). Note that this essentially consists in replacing *dν*(*t*) by *f* (*t*) *dt* in integrals, and that either of *f* and *𝓁* can be deduced from the other, since 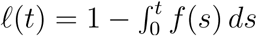 and *f* (*t*) = −*𝓁*^*′*^(*t*).

The age-specific fertility is the function *m*. If we denote by *M*_*i*_(*a, b*) the integer-valued random variable corresponding to the number of offspring produced by *i* between ages *a* and *b*, then assuming that *b* ⩽ *T*_*i*_ we have, as expected,

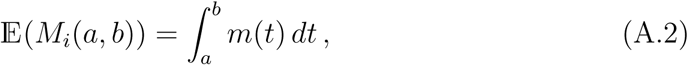

Obviously, the framework of Crump-Mode-Jagers processes is not meant to take into account all phenomena that shape the structure and dynamics of real-world populations. For instance, it assumes that individuals are independent and thus excludes any kind of density dependence. Similarly, the (optional) assumption that individuals reproduce at rate *m* is constraining, and in particular implies that they cannot produce several offspring simultaneously. Nevertheless, this framework is close to the minimal setting containing all the ingredients needed to define most descriptors of populations, whilst being simple enough to remain tractable and make it possible to derive explicit formulas for these descriptors. Moreover, the hypotheses above correspond quite well to the assumptions that are made, typically implicitly, to obtain the classic expressions of many of descriptors of populations.

Finally, to obtain discrete-time equivalents of formulas (1) and (3) we will need to consider the following version of the model, which allows simultaneous births: we keep assumption (i) under the extra hypothesis that the lifespan *T*_*i*_ is an integer-valued random variable, and we replace (ii) by the assumption that at each age *t* = 1, *…, T*_*i*_, individual *i* gives birth to 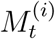 new individuals. Again, this corresponds quite well to the usual hypotheses on which many classic formulas rely.

## B The mean age of the parents of the offspring produced by a cohort

We now give a rigorous interpretation of the quantity *μ*_1_ given by formulas (1) and (2). As we will see, this interpretation is more subtle than what is usually assumed. This is because *μ*_1_ does *not* correspond to the expected value of the average of the ages of the parents of the offspring produced by a cohort, but only to the limit of this average when the size of this cohort goes to infinity.

Let 𝒞 denote a cohort, that is, a set of *n* individuals considered from the time of their birth to the time of their death. Let *T*_*i*_ be the lifespan of individual *i*, and *P*_*i*_ be the set of ages at which it produces offspring. Note that in our setting, conditional on *T*_*i*_, *P*_*i*_ is a point process with intensity *m* on [0, *T*_*i*_].

The average of the ages of the parents of the offspring produced by the cohort over its lifetime is

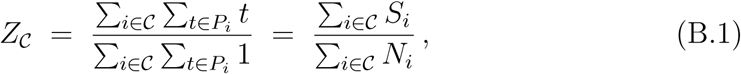

Where 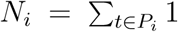 is the number of offspring produced by individual *i*, and 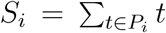 is the sum of the ages at which it produces them. Note that *Z*_C_ is well-defined only when Σ_*i∈𝒞*_ *N*_*i*_ *>* 0, but that this happens with probability arbitrarily close to one for a large enough cohort.

As we have already seen, the expected number of offspring produced by an individual *i* whose lifespan is *T*_*i*_ = *t* is

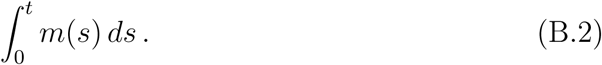

This quantity can be thought of as “𝔼(*N*_*i*_ *| T*_*i*_ = *t*)”, even though this interpretation is subject to some caution. At any rate, it follows that

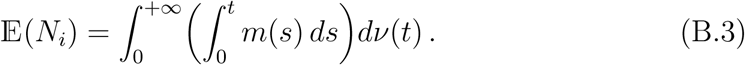

Moreover, using Fubini’s theorem,

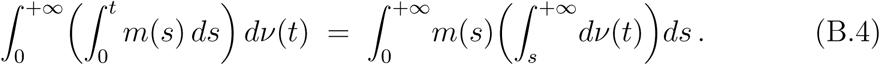

Using that 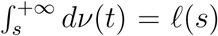, we get the well-known expression for *R*_0_, the *mean number of offspring produced by an individual during its lifetime*:

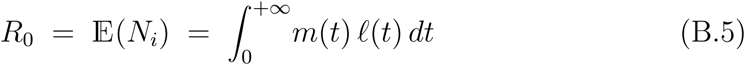

Using Campbell’s formula (equation (S1.11) in the Online Supplements) and the exact same reasoning, we can express the *mean sum of the ages at which an individual produces offspring* as

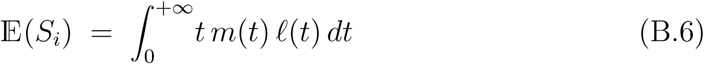

Let *N* (resp. *S*) denote a random variable that has the common distribution of the variables *N*_*i*_ (resp. *S*_*i*_). Then, as pointed out in most sources presenting the measure *μ*_1_, we have

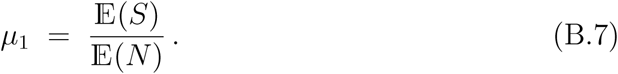

This however does not establish a link between *μ*_1_ and *Z*_*𝒞*_, the average age of the parents of offspring produced by the cohort. To see how these two quantities are related, observe that since the variables *N*_*i*_ (resp. *S*_*i*_) are independent, if we denote by *n* = Card(*𝒞*) the size of the cohort then by the law of large numbers, as *n →* +*∞*,

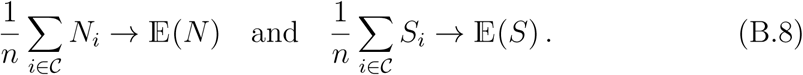

As a result,

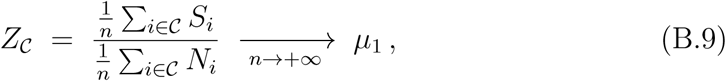

where the convergence is almost sure, i.e. happens with probability 1.

Importantly, note that *μ*_1_ is *not* the expected value of *S*_*i*_*/N*_*i*_ or of *Z*_*𝒞*_. In fact, the expected value of *S*_*i*_*/N*_*i*_ (conditional on this variable being well-defined) is precisely what we have termed the mean age at reproduction. We explain how to compute it in the next section.

## C The mean age at reproduction *τ*

Recall that we have defined the mean age at reproduction to be the expected value of the average of the ages at which a typical parent produces offspring. Formally, assuming that individual *i* has some offspring, the average age at which it produces them is

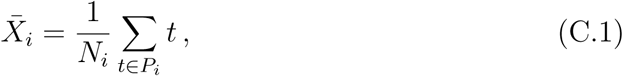

where, as before, *N*_*i*_ is the total number of offspring produced by *i* and *P*_*i*_ is the set of ages at which it produces them. The mean age at reproduction is thus

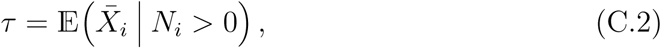

which, given our assumptions, does not depend on *i* or on the composition of the population.

To compute *τ*, let *I* be a “typical parent”, i.e. be uniformly sampled among the individuals that produce offspring during their lifetime. We then have

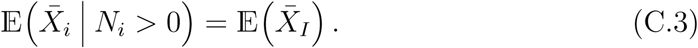

Moreover, letting 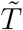 denote the lifespan of *I*, 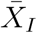 is the average of a point process with intensity *m* on 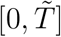. As explained in Section S1 of the Online Supplements, in the case of a Poisson point process, the expected value of this average is simply the expected value of a random point of 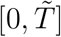 with density 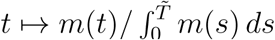. The remarkable fact that it does not depend on the value of *N*_*i*_ is a consequence of the absence of internal structure of Poisson point processes. From this, we get

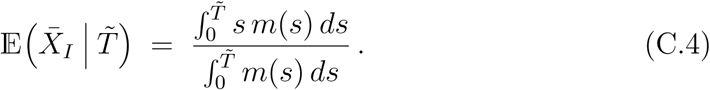

As a result,

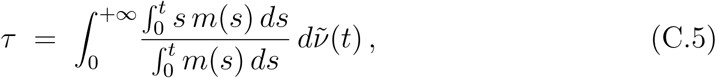

where 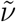 is the law of the lifespan 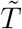 of *I*. Note that it is different from *ν*, the lifespan of a fixed individual, because conditioning on the fact that an individual produces offspring biases its lifespan; for instance, if – as frequently the case in real applications – there exists an age *α* such that *m*(*t*) = 0 for *t < α*, then individuals that produce offspring all live longer than *α*, whereas it is not necessarily the case for other individuals.

The last thing that we need to do in order to get an explicit formula for *τ* is thus to determine 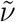. For this, note that

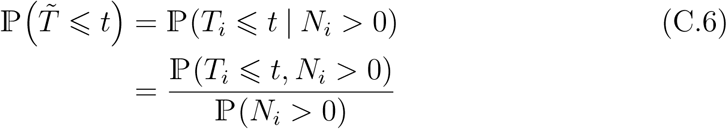

Conditioning on *T*_*i*_, using the void probabilities of Poisson point processes (see equation (S1.1) in the Online Supplements) for the probability that an individual with lifetime *s* produces some offspring, and finally integrating against *ν*, we get

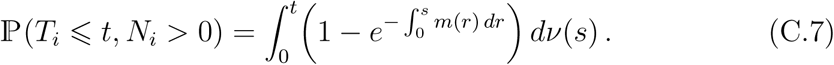

As a result,

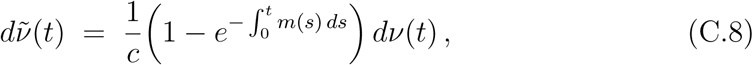

where the constant *c* =ℙ(*N*_*i*_ *>* 0) is given by

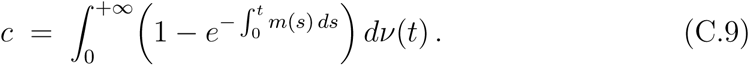

Note that, by integrating by parts and using that *𝓁*(*t*) *→* 0 as *t* → +∞, we can also express *c* directly in terms of *𝓁* and *m* as

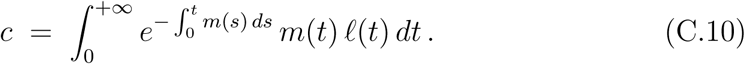

Putting the pieces together in the case where *T*_*i*_ has a density *f*, we get formula (1):

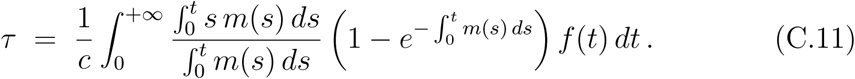

Note that neither the biological interpretation of *τ* nor the derivation of its expression depend on the assumption that individuals are independent.

Formula (10) for the average of a function *z* of the ages at which a parent produces offspring is obtained similarly, except that we have to work with

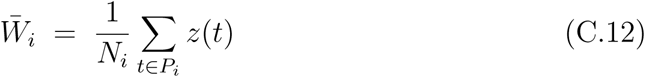

instead of 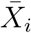, and use equation (S1.10) instead of equation (S1.6) to get

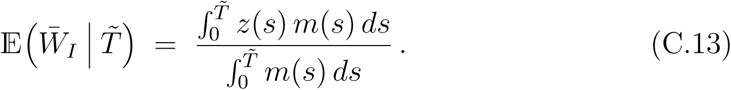

The justification of the expression of *τ* for discrete age structures can be found in Section S2 of the Online Supplements. It essentially consists in approaching the discrete-time model with the continuous-time one by choosing appropriate age-specific fertilities, and relies on the assumption that the number of offspring produced each year by each individual follows a Poisson distribution. It should also be pointed out that, because in the discrete-time setting individuals can produce several offspring simultaneously, there are two possibilities to define the average age at offspring production: counting all births equally, or weighting them by the number of offspring produced. Formula (2) is obtained by weighting the ages by the number of offspring produced when averaging them.

Finally, to obtain an equivalent of formula (5) for more general population structures, such as those allowed by matrix population models, one would need to (1) find the law of the conditional trajectory of an individual in the life cycle given that it produces offspring and (2) integrate the average of the ages at which it produces offspring against this law. While the first of these steps is feasible ^4^, it is unclear whether the resulting expression – if it can be obtained – would be simple enough to be useful.

## Online Supplements

### S1 Basic facts about Poisson point processes

In this section we recall basic results about Poisson point processes, focusing on the properties on which our calculations rely. Thus no attempt is made to state the results in full generality, and we do not preoccupy ourselves with technical conditions such as measurability. For a detailed presentation of Poisson point processes, see e.g. Kingman (1992) or Daley and Vere-Jones (2003).

It is common in modelling to assume that an event *occurs at rate r*(*t*) *at time t*. Loosely speaking, this means that the probability that the event happens between *t* and *t* + *dt* is independent of its previous occurrences, and is approximately *r*(*t*) *dt*. The rigorous way to formalize this is to say that the events are distributed according to an (inhomogeneous) Poisson point process with intensity *r*. Such a process can be seen as a random set of points characterized by the following properties: writing *N* (*I*) for the number of points that fall in a fixed set *I ⊂* ℝ,

i. *N* (*I*) is a Poisson random variable with mean ∫_*I*_ *r*(*t*) *dt*.
ii. *N* (*I*) and *N* (*J*) are independent whenever *I* and *J* are disjoint.

Note that the following useful fact is an immediate consequence of (i):

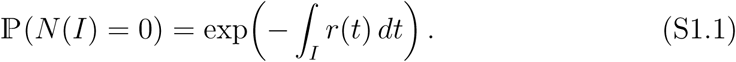

Property (ii), often known as the *independent scattering* property, essentially says that Poisson point processes have a “completely random” structure.

From now on, we consider a fixed set *I ⊂* ℝ such that *∫*_*I*_ *r*(*t*) *dt <* +∞. We let *P* be a Poisson point process with intensity *r* on *I* and denote by *N* = Card(*P*) its number of points. Let *X* be a random point of *I* with density *t* ↦ *r*(*t*)*/ ∫*_*I*_ *r*(*t*) *dt*, i.e. whose distribution is characterized by

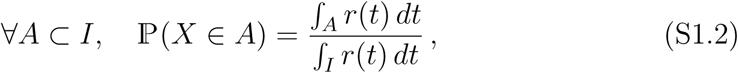

and note in passing that

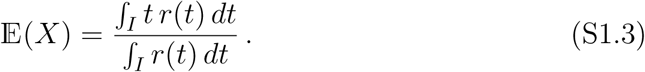

Then, conditional on *N* = *n, P* consists of *n* independent copies of *X* – that is, for every function *φ*,

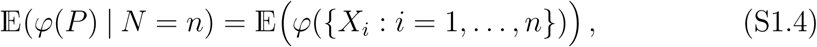

where *X*_*i*_, *i* = 1, *…, n*, are independent copies of *X*.

A consequence of this is that the expected value of the average of the points in *P* is 𝔼(*X*). Formally, if *N >* 0 then we can define a random variable 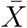 by

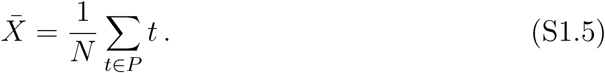

We then have

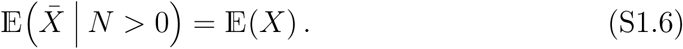

Indeed,

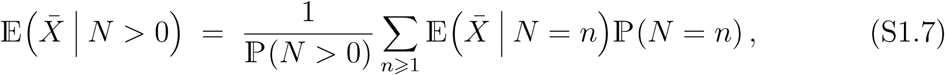

and, for every *n* ⩾ 1,

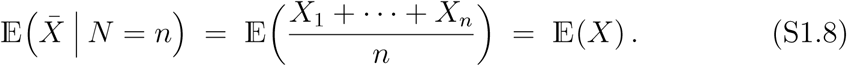

In fact, given a function *f*, the exact same reasoning can be applied to

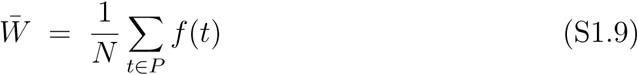

to show that

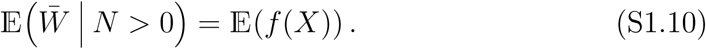

We close this short overview with a fundamental result known as Campbell’s formula. This formula states that, for every function *f*,

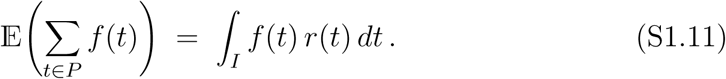

In contrast to (S1.6) and (S1.10), which are consequences of the independent scattering property, Campbell’s formula is not specific to Poisson point processes.

### S2 Expression of *τ* for discrete age structures

In discrete time, individual *i* has an integer-valued lifespan *T*_*i*_ and, at each age *t* = 1, *…, T*_*i*_, produces 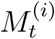 new individuals, where the variables 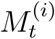 are integer-valued and independent of everything else. Here we will also need to assume that each variable 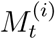 is a Poisson random variable with mean *m*_*t*_.

In that setting, the average 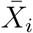 of the ages at which individual *i* produces offspring can be defined as

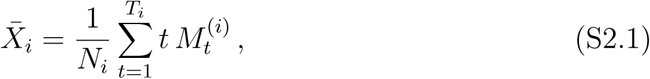

this definition being valid only when 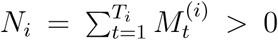 expression, each age at which *i* produces offspring is weighted by the number of offspring produced. This is similar to what is done for *μ*_1_, where each offspring contributes to the average age of the parents. But another possibility would be to weight all ages equally, that is, use the variable

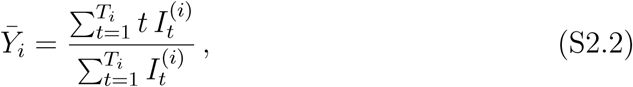

where 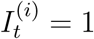 if 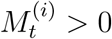 and 0 otherwise.

Since 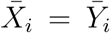 when individuals cannot give birth to several offspring simultaneously (or, more generally, when the number of offspring produced is either 0 or some constant *m*), the two definitions were equivalent in the continuous-time setting. But now, 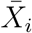 and 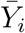 are two different and legitimate candidates for the “average age at which *i* produces offspring”. However, 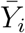 does not lend itself to analysis as easily as 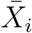 and to obtain formula (5) – which is arguably the natural discrete-time equivalent of formula (3) – it is 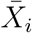 that should be used. Therefore, we define *τ* to be 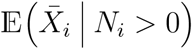.

The reasoning that lead to (3) could be adapted to obtain an expression for *τ*. However, it is also possible to deduce this expression directly from our results in continuous time. Indeed, the calculations of Section C of the Appendix are valid for general lifespans, including discrete ones: when *ν* is discrete, we simply have for any function *φ*

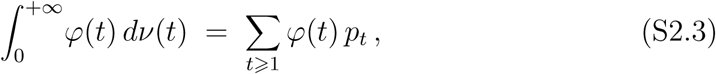

where *p*_*t*_ = ℙ(*T*_*i*_ = *t*).

Moreover, observe that if we let the age-specific fertility *m* be the piecewise constant function defined by

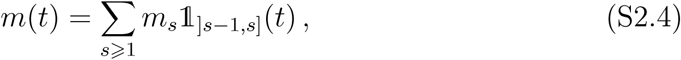

where 1_]*s -* 1,*s*]_ is the function that evaluates to 1 if *t* ∈]*s -* 1, *s*] and 0 otherwise, then the number of offspring produced by an individual between ages (*t -* 1) and *t* is a Poisson variable with parameter *m*_*t*_. Thus, the only difference with the discrete setting is that the ages at which these offspring are produced are uniformly distributed in] *t -* 1, *t*] instead of all equal to *t*.

Now, if we take the age-specific fertility to be the function *m*^(*ε*)^ defined by

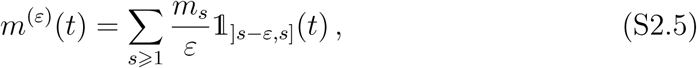

then the number of offspring produced between ages (*t -* 1) and *t* is still a Poisson variable with parameter *m*_*t*_, but this time the ages at which these offspring are produced are uniformly distributed in] *t - ε, t*]. Taking *ε* to zero, the mean age at childbirth will therefore tend to that of the discrete-time model. We spare the reader the straightforward but somewhat technical argument by which this can be made rigorous. Noting that, for continuous functions *g*,

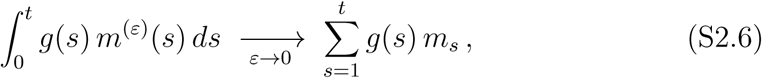

we obtain the following discrete-time equivalent of (3):

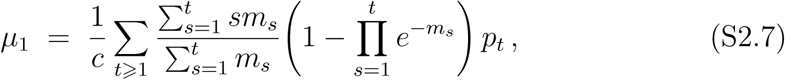

where *p*_*t*_ = ℙ(*T*_*i*_ = *t*) = *𝓁*_*t*_ - *𝓁*_*t*+1_, and

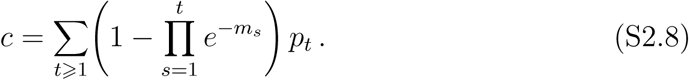

### S3 Proof of 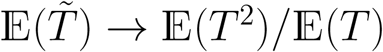 **as** *m →* 0

In this section we prove that, when the lifespan *T* of a fixed individual has a second moment – a condition that is always met in practice – and the age-specific fertility is constant and equal to *m*, then the expected lifespan of individuals that produce offspring during their lifetime converges to 𝔼(*T*^2^)*/* 𝔼(*T*) as *m →* 0. As seen in the main text, it follows immediately that *τ → μ*_1_, both in the continuous setting where offspring production occurs at a constant rate *m* during the lifetime of individuals and in the discrete setting where individuals produce a Poisson(*m*) number of offspring at each integer-valued age *t* ⩾ 1.

**Proposition 1.** *Let T denote the lifespan of a fixed individual, and let 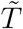 have the distribution of T conditional on reproduction in the model where reproduction happens at constant rate (or in the model where individuals produce* Poisson(*m*) *offspring at each integer age t*) 1 *at which they are alive), i.e.*

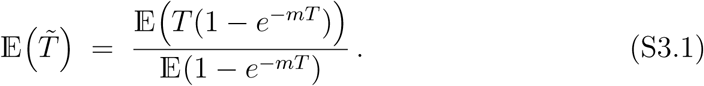

*Then, if* 𝔼(*T*^2^) *<* +*∞*,

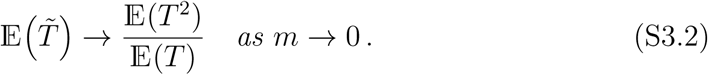

*Proof.* The following proof is due to Stephen P. Ellner and is a welcome simplification of my original proof.

Let *g*(*m, t*) = (1 - *e*^*-mt*^)*/m*, so that

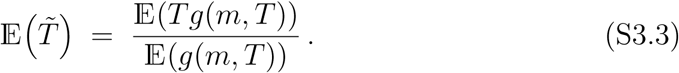

Since

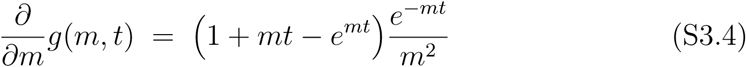

and that 1 + *x* ⩽ *e*^*x*^ for all *x*, we see that *g*(*m, t*) increases to *t* as *m* decreases to 0. By the monotone convergence theorem, it follows that 𝔼(*g*(*m, T*)) *↑*𝔼 (*T*) and 𝔼(*Tg*(*m, T*)) *↑*𝔼 (*T* ^2^) as *m ↓* 0. This terminates the proof.

### S4 Proof of *τ* ⩽ *μ*_1_

In this section we prove that *μ*_1_, as defined by formulas (1) and (2), is always greater than or equal to *τ*, as defined by formulas (3) and (5). This will be a simple consequence of the following lemma.

#### Lemma 1.

*Let X be a positive random variable, and let g and h be positive functions such that x* ↦ *g*(*x*)*/x is nondecreasing and x* ↦ *h*(*x*)*/x is nonincreasing. Then*,

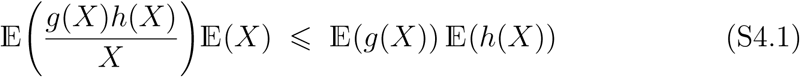

#### Proof

Let *Y* be a random variable with the same distribution as *X* and that is independent of *X*. We have to show

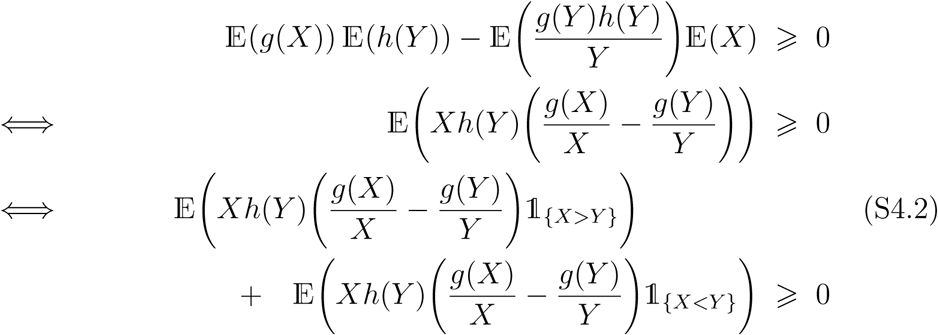

Since *x* ↦ *g*(*x*)*/x* is nondecreasing,

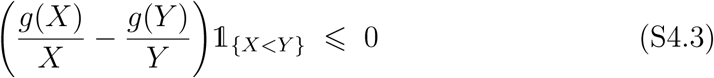

and since *x* ↦ *h*(*x*)*/x* is nonincreasing,

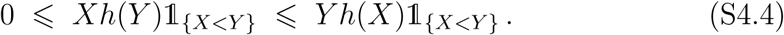

As a result,

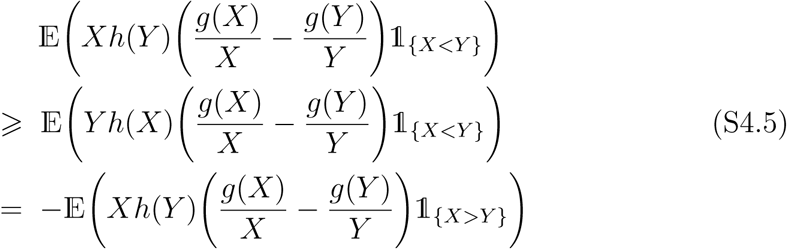

Plugging this into (S4.2) finishes the proof.

## Proposition 2

*Let T denote the lifespan of a fixed individual. Define the random variables M and M*^*∗*^ *by*

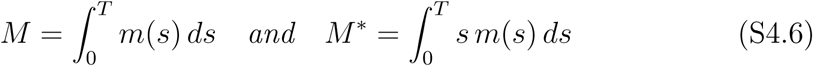

*in the case where reproduction occurs at a constant rate, and by*

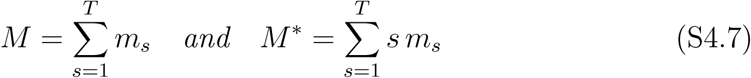

*in the case where it takes place at integer-valued ages t* ⩾ 1, *so that, in both cases*

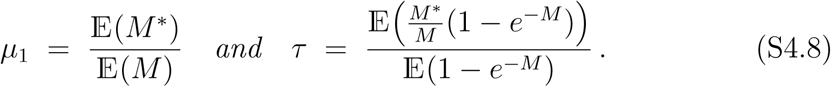

*Then, τ* ⩽ *μ*_1_.

### Proof.

First, observe that *M*^*∗*^ is actually a deterministic function of *M*. Indeed, let *ψ* (resp. *ψ*^*∗*^) denote the function such that *M* = *ψ*(*T*) (resp. *M*^*∗*^ = *ψ*^*∗*^(*T*)). Since *ψ* is nondecreasing, if we define *θ* by

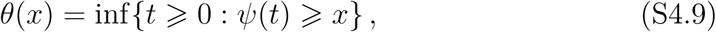

then we have *M*^*∗*^ = *ψ*^*∗*^(*θ*(*M*)). To see this, note that *θ*(*M*) ⩽ *T* by construction and that *θ*(*M*) *< T* implies 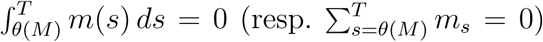, which in turn implies 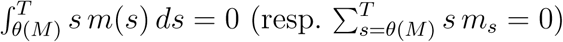. Thus, writing

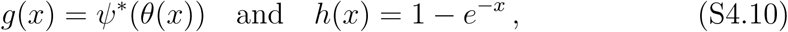

we have to prove

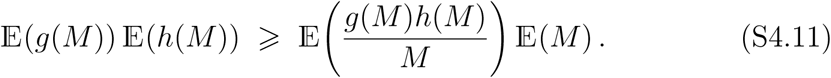

Clearly, *M* is a positive random variable, and the functions *g* and *h* are positive. Therefore, all we have to do to finish the proof is to show that *x* ↦ *h*(*x*)*/x* is nonincreasing and that *x* ↦ *g*(*x*)*/x* is nondecreasing, so that we can apply Lemma 1. First,

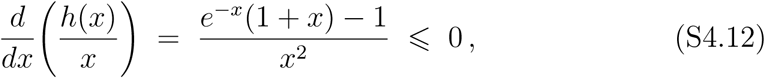

since 1 + *x* ⩽ *e*^*x*^. Second,

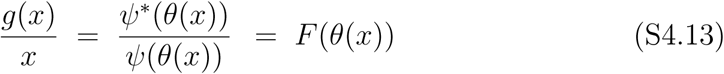

where *F*: *t* ↦ *ψ*^*∗*^(*t*)*/ψ*(*t*). The function *θ* is nondecreasing by construction. The fact that *F* is nondecreasing can be shown by straightforward calculations, e.g., in the continuous case,

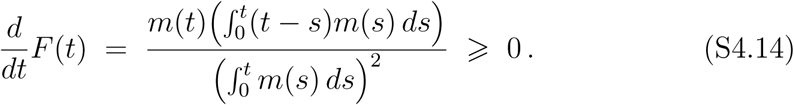

However, it is more satisfying to see that *F* (*t*) can be interpreted as the expectation of a random variable *X*_*t*_ with density *f*_*t*_(*s*) = *m*(*s*) 1 _[0,*t*]_(*s*)*/ψ*(*t*) in the continuous case, and with probability mass function 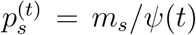 for *s* = 1, *…, t* in the discrete case. It then is easy to see that *X*_*t*_ is stochastically dominated by 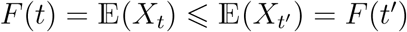 for *t < t*^*′*^, and so if follows immediately that *F* (*t*) = 𝔼(*X*_*t*_) 𝔼(*X*_*t*_*I*) = *F* (*t*^*′*^).

## S5 Example of calculation of *τ*

In this section we detail the calculations of *τ* in the case where the lifespan *T* of a fixed individual is a geometric variable such that for all *t* ⩾ 0, ℙ(*T*_*i*_ = *t*) = (1 - *p*)*p*^*t*^, and the age-specific fertility is constant and equal to *m* for all ages *t* ⩾ 1.

First, since

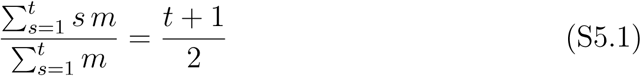

we see that, writing 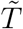 for the lifespan conditional on reproduction,

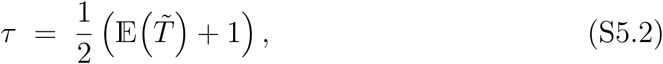

where

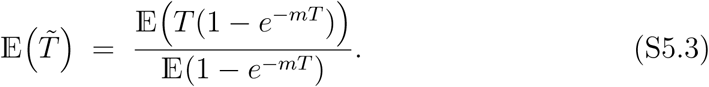

Now,

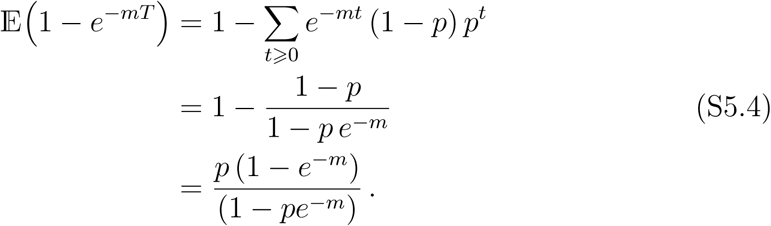

Similarly,

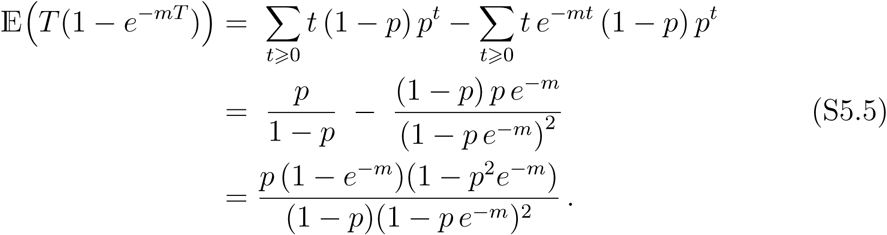

As a result,

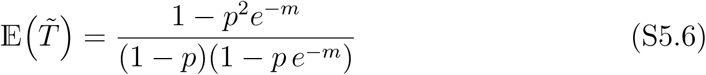

and finally,

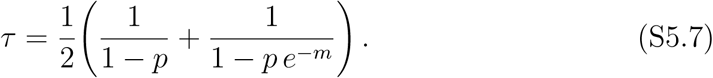

## S6 Projection matrices for *A. mexicanum*

In this section we give the projection matrices for *Astrocaryum mexicanum* that are behind Table 1. These matrices are taken from Appendix 6 of Cochran and Ellner (1992), who averaged them from several projection matrices of Pinero et al. (1984). Note that there is a small typo in the projection matrix for the *Low density* model given by Cochran and Ellner (1992): the entry (9, 8) of the projection matrix given is 0.8775, when it should be 0.08775. Correcting this, we find the same descriptors as in their Table 4.

The model is mostly size-based. Stage 1 corresponds to seedlings; stages 2–4 to non-reproducing juveniles and stages 5–10 to full-grown adults. In the matrices below, entries in bold correspond to reproductive transitions.

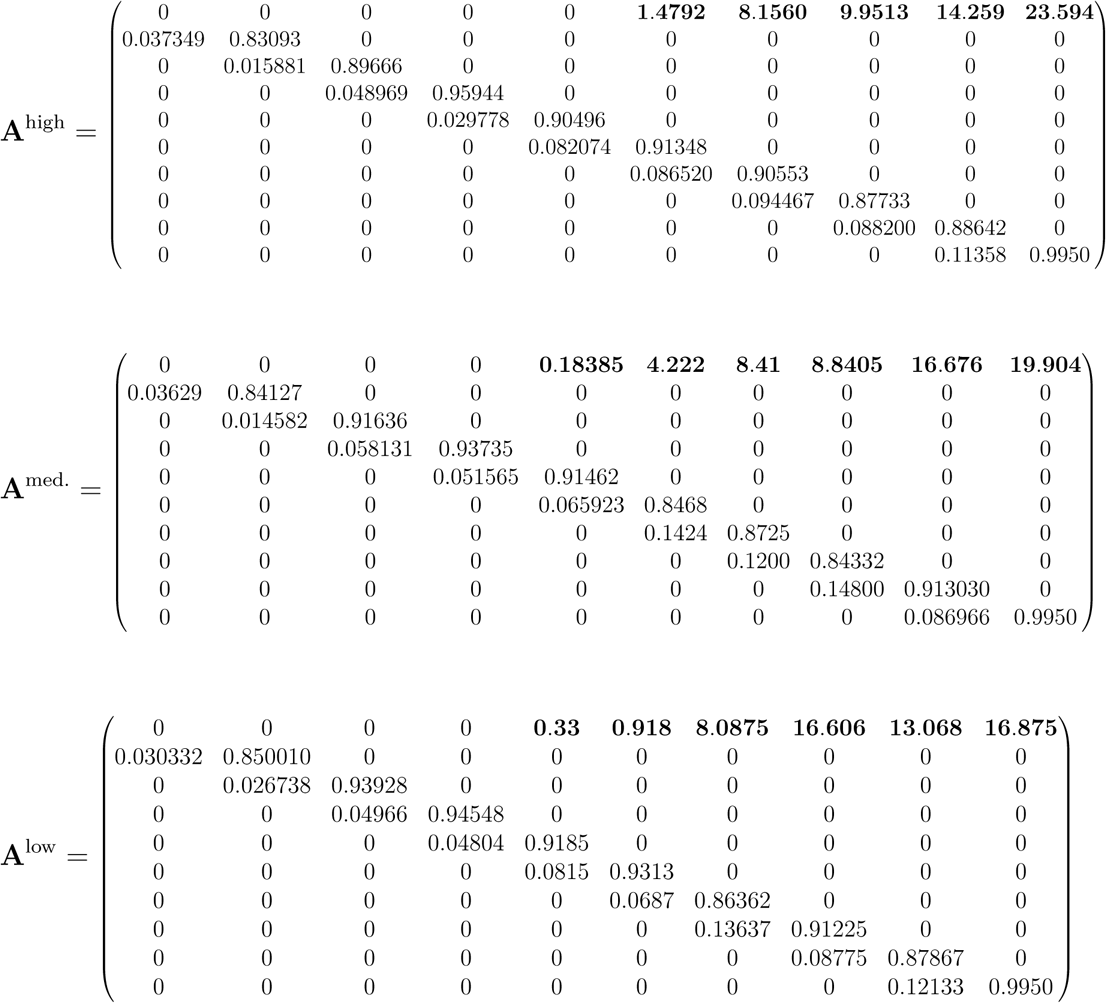

The estimates of *τ* and *L* (the expected lifespan conditional on reproduction) presented in Table 1 were obtained from these matrices using straightforward individual-based simulations. The code is available as an Online Enhancement to the manuscript.

## Footnotes

1 This was pointed out by Mauricio González-Forero.

2 What Keyfitz calls the *stationary population* is actually a cohort.

3 In probability theory and statistics, the *survival function* almost invariably refers to the complementary cumulative distribution function of *T*_*i*_, *t* ↦ ℙ(*T*_*i*_ *> t*). Here, however, we will stick to the convention used in biology.

4 This was explained to me by Stephen Ellner– see e.g. Chapter 3 of Ellner et al. (2016).

